# Imputation-Based Harmonization Mitigates Site Effects Without Data Leakage in Machine Learning Studies

**DOI:** 10.64898/2026.09.28.755204

**Authors:** Noah Hillman, Andrew A. Chen, Fengling Hu, Simon N. Vandekar, Randa Melhem, Lori L. Beason-Held, Theodore D. Satterthwaite, Christos Davatzikos, Haochang Shou, Russell T. Shinohara, Alzheimer’s Disease Neuroimaging Initiative (ADNI)

## Abstract

Neuroimaging studies that pool data across clinical sites often suffer from site effects — variability in imaging measures that arises from technical heterogeneity across sites as opposed to true biological signal. While numerous harmonization methods have been proposed to remove site effects from imaging features, less attention has been placed on how to incorporate harmonization models into common machine-learning pipelines. We demonstrate that current approaches for integration either suffer from data leakage, fail to fully remove site effects from the target data, or attenuate true biological associations during the harmonization process. To address these issues, we propose Multiple Imputation for Removing Technical Heterogeneity (MIRTH), which uses imputed outcomes to harmonize the test data. We benchmark MIRTH’s performance using both simulated and real-world volumetric data from the Alzheimer’s Disease Neuroimaging Initiative and the Baltimore Longitudinal Study of Aging, demonstrating that MIRTH can achieve high predictive accuracy while avoiding inflated performance when the outcome is imbalanced across clinical sites.

## 1 Introduction

Neuroimaging studies increasingly utilize data from large-scale multi-site cohorts to increase sample sizes and improve the reproducibility of reported findings (Dansereau et al., 2017; Kislik et al., 2025; Radua et al., 2020). While building these large multi-site data sets has facilitated scientific discovery across a wide range of research areas (Casey et al., 2018; Di Martino et al., 2014; Mueller et al., 2005; Thompson et al., 2014), pooling neuroimaging data across sites introduces unique analytical challenges. For example, it is well established that pooled neuroimaging data may contain systematic differences in imaging measurements across sites due to differences in scanner hardware, acquisition parameters, or research protocol (Han et al., 2006; Jovicich et al., 2009; Reig et al., 2009) — commonly referred to as *site* or *scanner* effects (Bayer et al., 2022). Even after attempts to standardize scanner hardware or acquisition parameters, inter-site differences in imaging features still commonly persist (Mirzaalian et al., 2016; Takao et al., 2011).

This problem has motivated data-driven harmonization methods that attempt to remove site effects directly from derived imaging features by jointly modeling biological and technical variability (Pinto et al., 2020; Tax et al., 2019; Yu et al., 2018). Hu et al. (2023) provide an overview of such methods, which run the gamut of Bayesian, latent variable, and deep learning approaches. One popular method, ComBat, uses an empirical Bayes linear model framework to remove the effect of site on both the mean and variance of imaging features while preserving associations with covariates of interest (Fortin et al., 2018; Johnson et al., 2007). The ComBat model has been validated on a wide variety of MRI studies (Fortin et al., 2017, 2018; Z. Li et al., 2026; Takahashi et al., 2026; Yu et al., 2018) and extended to accommodate more complex settings such as nonlinear associations (Pomponio et al., 2020), repeated measures within individuals (Beer et al., 2020), non-normal data (Horng et al., 2022), site effects on the covariance (Chen et al., 2022), and site-induced higher-order relationships between imaging features (Hu et al., 2024).

While ComBat and its extensions have been useful for removing site effects from neuroimaging data, less attention has been placed on how to integrate ComBat into common machine-learning workflows. Many studies have harmonized the entire neuroimaging data set prior to train-test splitting (Bourbonne et al., 2021; Campello et al., 2022; He et al., 2021; Ingalhalikar et al., 2021; Luna et al., 2021; Zhu et al., 2022), which may cause data leakage due to the harmonization model being fit and applied on the same sample. To fix this problem, researchers have proposed software that fits the harmonization model on the training data and transforms the test data by removing the batch effects learned on the training data (Marzi et al., 2024; Xin et al., 2025). Harmonization can therefore adhere to standard machine-learning principles that discourage jointly applying preprocessing steps to the training and test set (Kapoor & Narayanan, 2023).

Yet, a remaining challenge involves how to handle the outcome variable when performing image harmonization in predictive modeling contexts. When a trained prediction model is applied in real-world settings, the outcome will be unavailable and therefore unusable for harmonization. Typically, the researcher would want to include the outcome variable in the ComBat model so that imaging-outcome associations are preserved during harmonization. This is particularly important when the outcome is strongly imbalanced across sites, as ComBat removes residual inter-site differences in imaging measures and therefore could attenuate true imaging-outcome associations if the outcome is not included in the harmonization model. Thus, if the outcome is included in the harmonization model the researcher risks data leakage and limited applicability to real-world data settings; if the outcome is not included in the harmonization model then imaging-outcome associations become biased downwards when the outcome is imbalanced across sites.

Nieto et al. (2026) proposed to address this problem by using multiple sets of surrogate outcome labels to create harmonized data sets from which an ensemble model can generate a final prediction. While this approach exhibited strong performance across a wide range of prediction tasks, creating pretend outcome values for continuous outcomes involves discretization that could limit model flexibility and impact performance. Additionally, using surrogate outcomes does not accommodate more complex settings where missing data are present in both covariate and outcome variables. Therefore, it remains to be seen whether standard approaches for missing data (Dong & Peng, 2013) could widen the applicability of current methods while also maintaining high predictive performance.

One popular framework for handling missing data is multiple imputation (MI), where a small number of complete data sets are generated by replacing missing data with plausible values generated from conditional models fit on the observed data (Austin et al., 2021; P. Li et al., 2015; Rässler et al., 2013). MI improves upon single imputation by propagating uncertainty in the imputation model into downstream inferential and prediction tasks, thereby improving standard error estimates and potentially mitigating bias (Murray, 2018; Rubin, 1987). Moreover, current software for implementing MI (van Buuren & Groothuis-Oudshoorn, 2011) allows researchers to easily generate multiple imputed data sets even when numerous variables contain missing values.

In this paper, we propose Multiple Imputation for Removing Technical Heterogeneity (MIRTH) for integrating ComBat into machine-learning pipelines. Similar to other approaches, MIRTH aims to prevent data leakage by estimating batch effect parameters on the training data set and applying these estimates when removing site effects from the held-out test data. Furthermore, MIRTH attempts to preserve imaging-outcome associations during the harmonization process by using imputed outcomes (and potentially other covariates) to accommodate missing data present in the test set. We benchmark the performance of MIRTH across a wide range of potential data settings through simulation studies. Additionally, we demonstrate MIRTH’s utility in real data applications using volumetric data from the Alzheimer’s Disease Neuroimaging Initiative (Mueller et al., 2005; Petersen et al., 2010) and the Baltimore Longitudinal Study of Aging (Ferrucci, 2008; Shock et al., 1984) to predict age, sex, and Alzheimer’s disease.

## 2 Methods

### 2.1 MIRTH Overview

Let 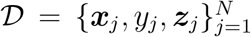 denote a neuroimaging data set, where each participant *j ∈ {*1, …, *N }* is scanned at a site *i ∈ {*1, …, *S}*, ***x***_*j*_ *∈* ℝ^*V*^ is a *V* -dimensional vector of derived imaging features (e.g., volumetric measures at each region of interest), *y*_*j*_ *∈ Y* is the outcome of interest with *Y ⊆* ℝ for continuous outcomes or *Y* = *{*0, …, *C ™* 1*}* for categorical outcomes with *C* classes, and ***z***_*j*_ is a *Q*-dimensional vector of non-imaging covariates (e.g., demographic or clinical variables). Partition *D* into training and test subsets such that

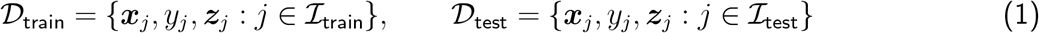

where the same sites appear in the training and test set, *ℐ*_train_ *∪ℐ*_test_ = *{*1, …, *N }, ℐ*_train_ *∩ℐ*_test_ = *∅*, and*n*_tr_ = |*I*_train_|. To ensure applicability in real-world data settings where the outcome is unavailable, the test outcomes *{y*_*j*_ : *j ∈ ℐ*_test_*}* are treated as missing. While presently only covariates and outcomes contain missing values, MIRTH could be extended to settings where the imaging data also contain missingness.

MIRTH, which is outlined in Figure 1, addresses missing data through the following procedure:

**Figure 1:**
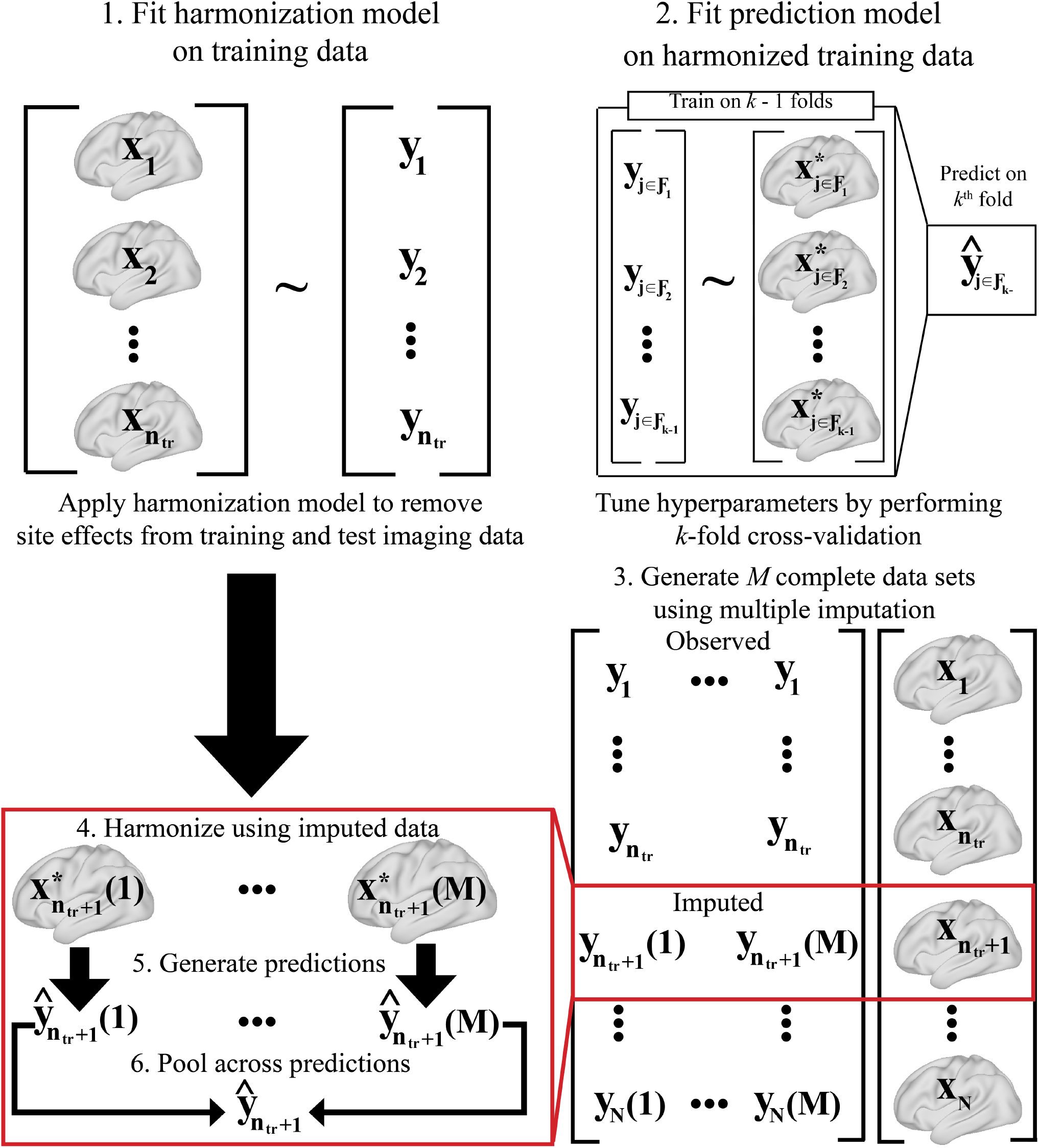
An overview of MIRTH, an algorithmic procedure for integrating ComBat into common machine-learning workflows. First, the ComBat harmonization model is fit on the training data set. Next, a prediction model is fit on the harmonized training data, with hyperparameters tuned through a *k*-fold cross-validation procedure. Afterwards, *M* imputed test data sets are generated via multiple imputation. While in the figure only the outcome is imputed, the procedure can be easily generalized to other missing covariates. Using the fitted harmonization model from step 1, each of the imputed test data sets is harmonized. After applying the prediction model from step 2 to get intermediate predictions for each imputed data set, predictions are pooled across imputations to get a final prediction for each individual.

1. First, fit and apply the ComBat model on *D*_train_. While ComBat has been described in further detail elsewhere (Fortin et al., 2017; Johnson et al., 2007), using the notation above ComBat assumes that for a participant *j* scanned at site *i*, the observed value of feature *v* arises from the linear model

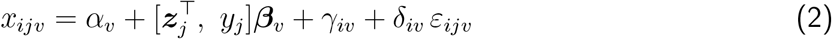

where *α*_*v*_ is the feature-level intercept, ***β***_*v*_ *∈* ℝ^*Q*+1^ captures associations with non-imaging covariates, *γ*_*iv*_ and *δ*_*iv*_ are the respective additive and multiplicative scanner effects, and *ε*_*ijv*_ *∼ N* (0, 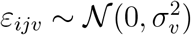) captures the error distribution. ComBat uses an empirical Bayes framework to estimate the model parameters, with the harmonized training data represented as

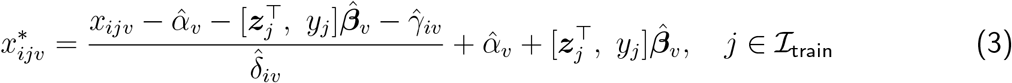

Let 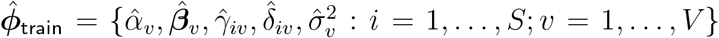 denote the full set of parameters estimated by the ComBat model fit on the training data set. Then, by denoting the harmonization operator described in (3) as *ℋ* 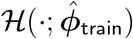, the harmonized training data can be represented as

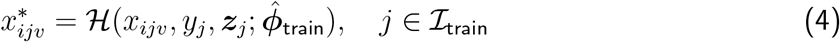

The complete training data set after harmonization is represented by 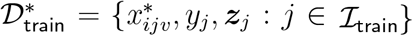.
2. Next, let *f*_***θ***_ : R^*V* +*Q*^ *→ Y* denote a prediction model indexed by parameters ***θ*** *∈* Θ with a vector of hyperparameters ***λ*** *∈* Λ. The model *f*_***θ***_ can be selected by the researcher to fit the underlying prediction task. Let 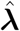 represent the set of hyperparameters that minimizes the out-of-fold prediction error when performing a *k*-fold cross-validation procedure on the training set. Using 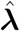, a final prediction model is fit on the entire harmonized training data set, which we denote 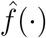.
3. To account for the missing data present in the test set, perform multiple imputation by chained equations (MICE) by fitting an imputation model on the unharmonized training data and generating *M* imputed test data sets (Raghunathan et al., 2001; van Buuren, 2007; van Buuren & Groothuis-Oudshoorn, 2011). MICE cycles through each variable with missingness, drawing from the posterior predictive distribution to generate a new imputation based on the observed and previously imputed values of other variables. While the imputation model can be chosen by the researcher, it should be congenial with the final prediction model (Murray, 2018). The algorithm typically stabilizes after a small number of cycles, yielding *M* imputed test sets 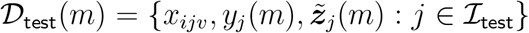 or each *m ∈ {*1, …, *M }* where

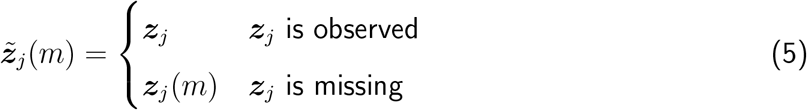

and *y*_*j*_(*m*), ***z***_*j*_(*m*) denote the *m*^th^ imputed value of each respective variable.
4. For each imputed test data set, apply the harmonization operator to yield

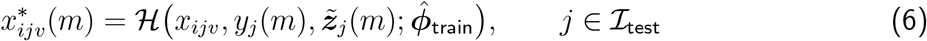

Note that MIRTH prevents data leakage by using batch effect parameters estimated on the training data when harmonizing the test set.
5. Apply the model 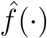 to get predictions 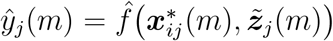 for each participant *j ∈ ℐ*_test_ and harmonized test set *m ∈ {*1, …, *M }*.
6. Pool across the *M* predictions to get a final prediction

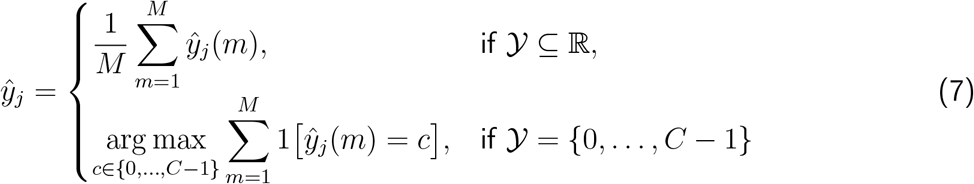

For categorical outcomes, one can also average across the predicted probabilities from each imputed data set to obtain a final probabilistic prediction.

### 2.2 Comparison Methods

MIRTH was evaluated against four potential harmonization procedures for building predictive algorithms on imaging features.

1. **No Harmonization:** Fit and apply a prediction model on unharmonized data, such that site effects are not removed from the training or test set.
2. **Oracle:** Fit the ComBat harmonization model on the training set and use the true test outcomes when applying the fitted model to harmonize the test set. Although Oracle suffers from data leakage (Nieto et al., 2026) and is not widely applicable — the true outcome values would be missing in real-world settings — it establishes a useful benchmark for MIRTH performance.
3. **No Outcome:** Fit the ComBat model on the training set while excluding the outcome from the harmonization model. Then, use the parameters estimated by the fitted model to harmonize the test set.
4. **PrettYharmonize:** As proposed by Nieto et al. (2026), first harmonize the training set using ComBat and fit a prediction model on the harmonized data. Next, generate a harmonized test data set by applying the fitted harmonization model to the test data while pretending every participant has the same outcome value. Repeat this procedure for each class (categorical response) or every decile (continuous response), such that intermediate predictions can be generated by applying the prediction model to each harmonized test data set. Train and apply a stack model that takes as input the predictions from each harmonized test data set to generate a final prediction. PrettYharmonize was implemented with publicly available software (https://github.com/juaml/PrettYharmonize) while using the random forest algorithm for the prediction model and either a logistic regression (categorical outcome) or random forest (continuous outcome) for the stack model.

### 2.3 Data

To validate MIRTH, a series of predictive tasks were conducted using structural MRI data from the Alzheimer’s Disease Neuroimaging Initiative (ADNI) and the Baltimore Longitudinal Study of Aging (BLSA), aggregated through the Imaging-based coordinate SysTem for Aging and NeurodeGenerative diseases (iSTAGING) consortium (Habes et al., 2021). Details on the image acquisition protocols for the ADNI and BLSA cohorts have been published elsewhere (Jack Jr. et al., 2008, 2024; Resnick et al., 2000, 2003). Participants’ T1-weighted (T1w) images underwent standard preprocessing steps including bias correction for intensity inhomogeneities (Tustison et al., 2010) and skull stripping (Doshi et al., 2013). T1w images were segmented into 145 regions of interest (ROIs) using a multi-atlas ensemble approach (Doshi et al., 2016) that has exhibited strong performance (Asman et al., 2013) by capturing uncertainty in the registration process when fusing atlas labels for a final segmentation. Volumetric measures were captured for each of the 145 ROIs that span both subcortical and cortical regions.

The ADNI + BLSA data set was subset to include the most recent visit from cognitively normal (CN) and Alzheimer’s disease (AD) participants with observed demographic, imaging, and diagnostic data. Table 1 provides an overview of the sample characteristics after sub-sampling, which intentionally created a cross-sectional sample that prioritized older ages where classification may become more difficult. For the ADNI cohort, participants were classified into CN and AD groups based on self-reported memory complaints as well as clinical assessments that took into account performance on the Wechsler Logical Memory II paragraph recall test (Wechsler, 1987), Clinical Dementia Rating (Morris, 1993), and Mini-Mental State Examination (Folstein et al., 1975), as described by the ADNI clinical core (Aisen et al., 2010; Petersen et al., 2010). Participants from the BLSA cohort were classified based on previously defined criteria (Wanigatunga et al., 2022), where participants were flagged for review if their Clinical Dementia Rating was *≥* 0.5 or they committed four or more errors on the Blessed Information-Memory-Concentration Test. AD designations were made based on diagnostic criteria laid out in the Diagnostic and Statistical Manual of Mental Disorders (American Psychiatric Association, 1994) and the National Institute of Neurological and Communicative Disorders and Stroke – Alzheimer’s Disease and Related Disorders Association (McKhann et al., 1984).

**Table 1.** Participant characteristics by scanner manufacturer and field strength, which are combined to create the site variable. Continuous variables are summarized as mean (SD) while categorical variables are reported as N (%). CN = cognitively normal and AD = Alzheimer’s disease.

|  | GE |  | Philips |  | Siemens |  |
| --- | --- | --- | --- | --- | --- | --- |
|  | 1.5T<br>(N = 285) | 3T<br>(N = 249) | 1.5T<br>(N = 48) | 3T<br>(N = 1,138) | 1.5T<br>(N = 218) | 3T<br>(N = 629) |
| **Age** | 77.8 (7.9) | 76.0 (7.4) | 76.9 (6.2) | 69.5 (15.0) | 78.8 (6.5) | 74.9 (7.8) |
| **Sex** |  |  |  |  |  |  |
| Female | 120 (42) | 118 (47) | 15 (31) | 623 (55) | 100 (46) | 349 (55) |
| Male | 165 (58) | 131 (53) | 33 (69) | 515 (45) | 118 (54) | 280 (45) |
| **Diagnosis** |  |  |  |  |  |  |
| CN | 126 (44) | 138 (55) | 12 (25) | 1,042 (92) | 65 (30) | 404 (64) |
| AD | 159 (56) | 111 (45) | 36 (75) | 96 (8) | 153 (70) | 225 (36) |

### 2.4 Simulation Study

To evaluate MIRTH across a range of potential data settings, we first conducted a simulation study where we systematically varied the sample size and “disease” effect magnitude. Simulated data were generated according to the ComBat model where for participant *j* at site *i* and imaging feature *v*, the simulated imaging measurement *x*_*ijv*_ was generated as

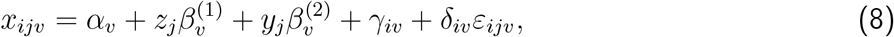

where *α*_*v*_ is the ROI-specific intercept, *z*_*j*_ is a nuisance covariate, *y*_*j*_ is the outcome of scientific interest, *γ*_*iv*_ and *δ*_*iv*_ are the additive and multiplicative site effects, and 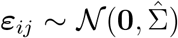 where 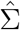 is the empirical correlation matrix of the imaging features in the ADNI + BLSA data set after regressing out age, sex, and AD diagnosis. The indices range over sites *i* = 1, …, *S*, participants *j* = 1, …, *N*, and ROIs *v* = 1, …, 145 such that the dimensionality of the feature space resembles the ADNI + BLSA data set. The mean additive and multiplicative site effect magnitudes were uniformly spaced across sites, such that given *π*_*r*_, the *r*^th^ random permutation of *{*1, …, *S}* where *π*_*r*_*{i}* is the *i*^th^ element of the shuffled vector, the site effect parameters are sampled from

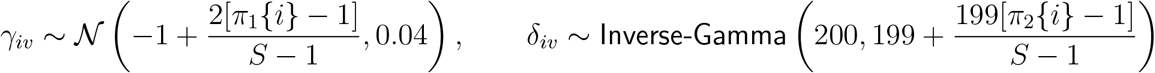

In other words, site-effect parameters are assumed to have uniformly spaced means from [*™*1, 1] and [1, 2] for location and scale batch effects respectively, while also allowing site effects to vary across features. The total sample size (*N ∈ {*250, 1000*}*) was varied across simulations while setting *S* = 5.

For non-imaging variables, *z*_*j*_ *∼ N* (0, 1) denotes a nuisance covariate while *y*_*j*_ represents the variable of scientific interest (i.e., the response variable in the prediction model). All imaging features and the nuisance covariate are included in both the imputation and prediction models. To examine performance when predicting both categorical and continuous outcomes, *y*_*j*_ was generated under two different regimes. For a binary classification setting, we assumed that *y*_*j*_ *∼* Bernoulli(*p*_*i*_) with a site-specific prevalence 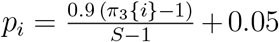, which corresponds to uniform spacing from [0.05, 0.95] across sites. To emulate predicting a continuous variable, we let *y*_*j*_ *∼ N* (0, 1) while ensuring that the polyserial association between the site and outcome variables is 0.75. Thus, in both binary and continuous settings we simulate data such that *y*_*j*_ is correlated with site.

For both binary and continuous outcomes, we explored null settings where 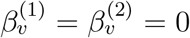 across all *v*, such that the imaging features were not predictive of the outcome. Note that since the outcome is correlated with site, poor harmonization could lead to above chance predictive performance even when no imaging-outcome association exists. Additionally, we conducted simulations under the alternative where 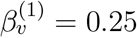 and

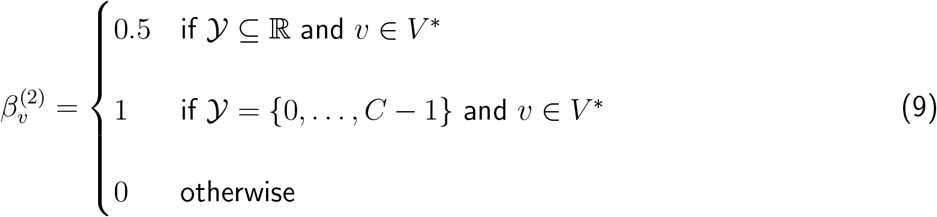

where *V* ^*\**^ is a random subset of imaging features such that |*V* ^*\**^| = 10.

To evaluate how MIRTH performed in settings where missingness is present in both outcome and covariate variables, *z*_*j*_ was randomly masked in the test set using the model Pr(*R*_*j*_ = 1) = 1*™*expit(*™*2+*y*_*j*_) where *R*_*j*_ is an indicator variable that denotes whether *z*_*j*_ is observed. We conducted these simulations using the aforementioned alternative setting to generate the simulated data, such that a subset of imaging features were associated with the outcome. Note that predictive performance under fully observed and missing covariate settings can be compared directly using the same simulated data by masking simulated covariate values.

Altogether, for each combination of sample size, outcome type (binary or continuous), association strength, harmonization approach, and missingness mechanism, 100 plasmode simulations were run. Simulated data were randomly split into a training and testing set (80% training), with each harmonization approach evaluated based on predictive performance in the test set. For MIRTH, a random forest imputation model was used with *M* = 25 imputations. A random forest model with *B* = 500 trees was used as the predictive model, with the number of variables to consider when splitting at each node *q*^*′*^ *∈ {*1, 3, …, 25*}* tuned via a 5-fold cross-validation procedure performed on the training set. To assess whether the harmonized test data contained residual site effects, random forest models were trained to predict site after regressing out covariate effects. In this setting, multi-class AUC (Hand & Till, 2001) values close to 0.5 indicated superior removal of site effects. Additionally, likelihood ratio tests were conducted to examine whether a full model with site and the true outcome value explained significantly more variance in the predicted outcome than a reduced model with only the true outcome value as a covariate, with significant associations suggesting that site effects were not fully removed from the test set prior to prediction.

### 2.5 Data Analysis

After conducting simulation studies, MIRTH was evaluated on the previously described ADNI + BLSA data set. Similar to the plasmode simulations, random forest models with *B* = 500 trees and hyperparameter tuning via 5-fold cross-validation were trained on 80% of the ADNI + BLSA data and applied to the test set, which was repeated 100 times to ensure that model results did not depend on the train-test split. Multiple imputation was performed using the mice R package (van Buuren & Groothuis-Oudshoorn, 2011) with a random forest model used to generate *M* = 25 imputations.

To examine how MIRTH performed across multiple outcome types, volumetric data from 145 ROIs were used to predict age, sex, and AD diagnosis. The two aforementioned variables that were not the outcome for a given prediction task were included as covariates alongside imaging features when fitting imputation and prediction models. Predictive performance was evaluated primarily using area under the receiver operating characteristic curve (AUC) for categorical responses and the root mean squared error (RMSE) for continuous outcomes. The effectiveness of the harmonization procedure was also evaluated by fitting random forest models to harmonized test data to predict site, where site is defined as all pairwise combinations of scanner manufacturer (GE, Siemens, Philips) and field strength (1.5T or 3T) similar to previous studies (Beer et al., 2020; Chen et al., 2022; Hu et al., 2024). For harmonization, age, age^2^, sex, and AD diagnosis were included in the ComBat model to preserve biological associations while removing technical variability across sites.

## 3 Results

### 3.1 Simulation Study

Figure 2 presents results from the null simulation setting where there is no association between simulated imaging values and the binary or continuous outcome of interest. Since the outcome is imbalanced across sites, poor harmonization could lead to inflated performance due to the predictive algorithm leveraging site effects to generate more accurate predictions. Successful harmonization pipelines should lead to chance prediction in this simulation setting, which corresponds to AUC = 0.5 for binary outcomes and RMSE = 1 for continuous outcomes. For binary outcomes and *N* = 1000, only the No Harmonization approach led to markedly optimistic predictions (AUC_median_ = 0.74), with all other methods exhibiting similar performance (AUC_median_ = 0.52–0.55). However, for continuous outcomes and *N* = 1000, both PrettYharmonize and No Harmonization exhibited inflated performance, with median RMSE values of 0.88 and 0.81, respectively.

**Figure 2:**
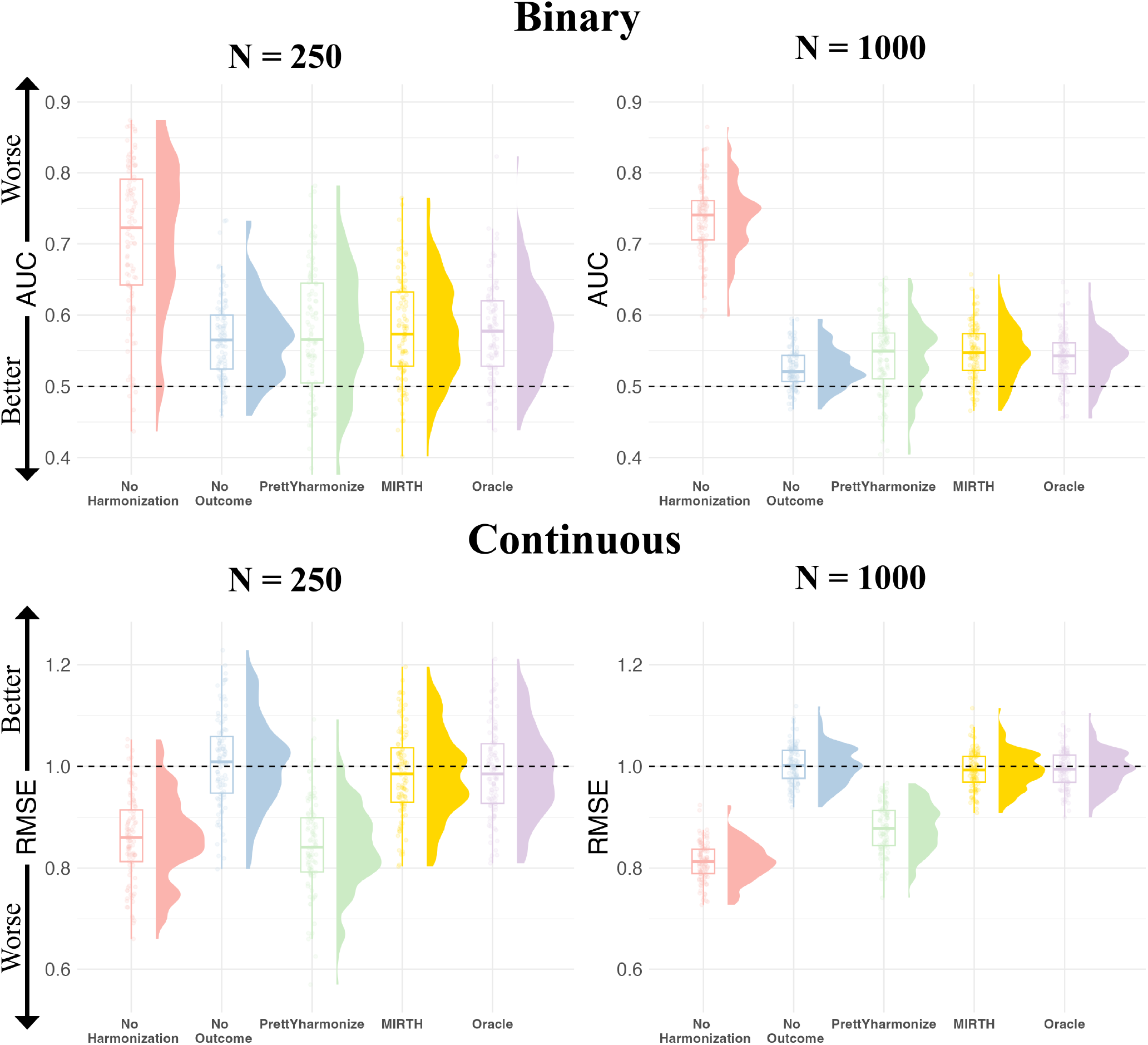
Predictive performance for null simulation settings where outcomes are independent of imaging features but correlated with site, such that poor harmonization can lead to above chance performance. Evaluation metrics are aggregated across 100 simulations with an 80*/*20 train-test split to evaluate whether a given harmonization procedure leads to inflated performance in the test set for either binary or continuous outcomes. Results are depicted for *N ∈ {*250, 1000*}* and *S* = 5.

We further evaluated whether each harmonization method removed site effects from simulated data when a true imaging-outcome association was present at |*V* ^*\**^| = 10 features (see Section 2.4 for details). Table 2 contains results from likelihood ratio tests that suggest that when the outcome is continuous both No Harmonization and PrettYharmonize generate predictions that are associated with site after controlling for the true outcome value (*p*_median_ *<* 0.001 for both methods). For continuous outcomes, only MIRTH and Oracle generated site-independent predictions across all sample sizes (*p*_median_ = 0.35–0.52). Moreover, when applying random forest models to predict site using the harmonized test data, MIRTH and Oracle exhibited null performance across sample size and outcome types with AUC_median_ = 0.47–0.52. Figure 3 presents model performance when predicting the outcome under the same simulation setting where *S* = 5 and |*V* ^*\**^| = 10 imaging features are associated with the outcome. Across all sample sizes and outcome types, No Outcome led to the worst prediction performance (AUC_median_ = 0.73–0.76; RMSE_median_ = 0.95–0.96). For binary outcomes, PrettYharmonize, MIRTH, and Oracle all exhibited comparable performance regardless of sample size, with ΔAUC *≤* 0.02 for all pairwise comparisons between the three models. However, for continuous outcomes, RMSE was markedly lower for Pret-tYharmonize (RMSE_median_ = 0.66–0.73) relative to MIRTH (RMSE_median_ = 0.83–0.86) and Oracle (RMSE_median_ = 0.84–0.87). The decrease in RMSE for PrettYharmonize and strong predictive performance of the No Harmonization approach corresponded with site-correlated predictions as shown in Table 2, such that in settings where the outcome is correlated with site, high predictive accuracy may result from a failure to remove site effects.

**Table 2.**
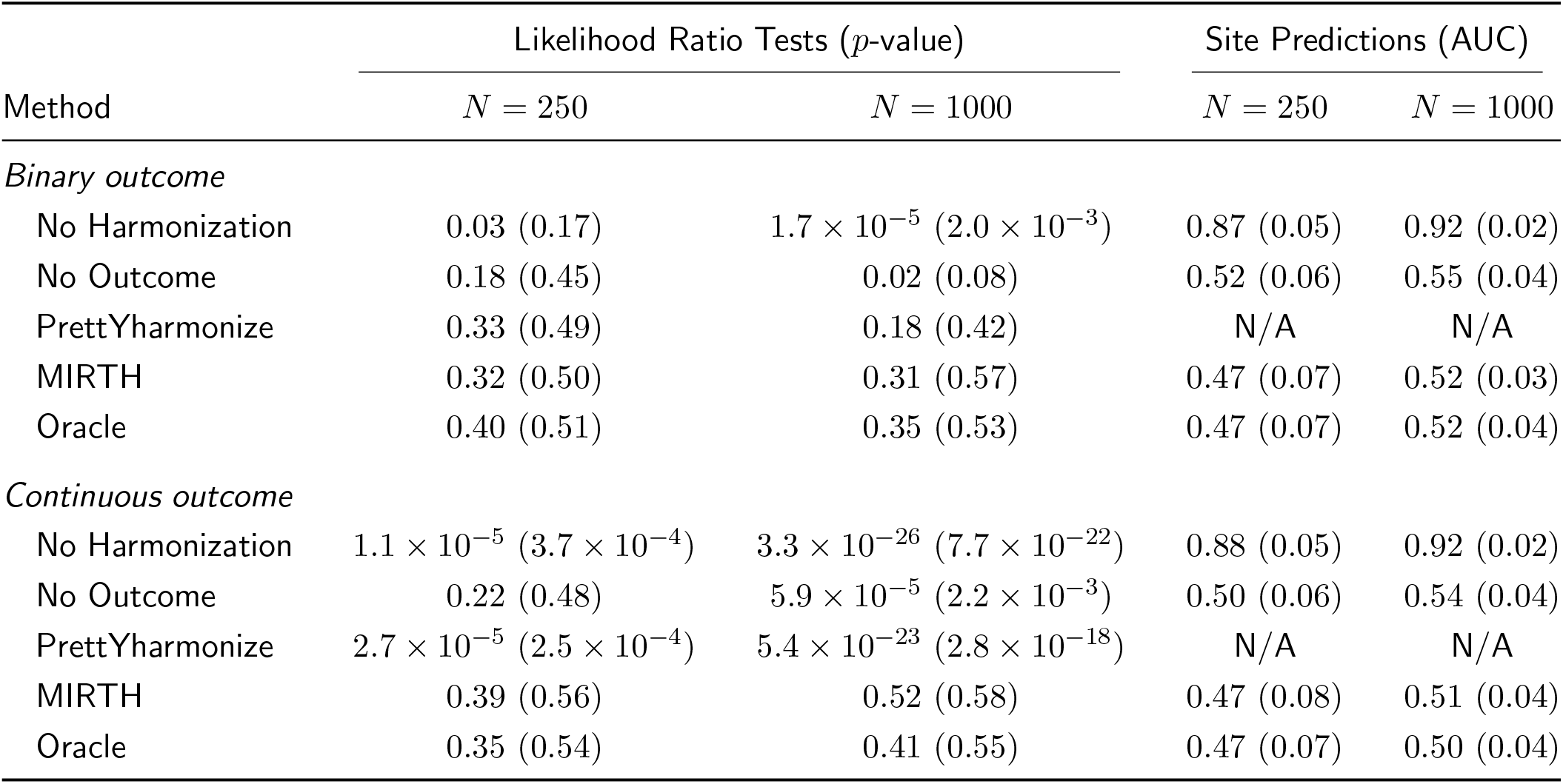
Median (IQR) values across 100 simulations (5 sites and nonzero imaging-outcome associations) for binary and continuous outcomes and two sample sizes, examining whether each harmonization method successfully removes site effects. The *p*-values are derived from likelihood ratio tests exploring whether outcome predictions in the test set are associated with site after controlling for the true outcome. Random forest models were also trained to predict site in the test set, with predictive performance evaluated via a multi-class AUC criterion (Hand & Till, 2001). Note that since PrettYharmonize does not attempt to generate a unique harmonized test data set that is independent of site, we do not evaluate PrettYharmonize along this dimension.

| Method | Likelihood Ratio Tests ($p$-value) | | Site Predictions (AUC) | |
| --- | --- | --- | --- | --- |
| | $N = 250$ | $N = 1000$ | $N = 250$ | $N = 1000$ |
| *Binary outcome* |  |  |  |  |
| No Harmonization | 0.03 (0.17) | $1.7 \times 10^{-5}$ ($2.0 \times 10^{-3}$) | 0.87 (0.05) | 0.92 (0.02) |
| No Outcome | 0.18 (0.45) | 0.02 (0.08) | 0.52 (0.06) | 0.55 (0.04) |
| PrettYharmonize | 0.33 (0.49) | 0.18 (0.42) | N/A | N/A |
| MIRTH | 0.32 (0.50) | 0.31 (0.57) | 0.47 (0.07) | 0.52 (0.03) |
| Oracle | 0.40 (0.51) | 0.35 (0.53) | 0.47 (0.07) | 0.52 (0.04) |
| *Continuous outcome* |  |  |  |  |
| No Harmonization | $1.1 \times 10^{-5}$ ($3.7 \times 10^{-4}$) | $3.3 \times 10^{-26}$ ($7.7 \times 10^{-22}$) | 0.88 (0.05) | 0.92 (0.02) |
| No Outcome | 0.22 (0.48) | $5.9 \times 10^{-5}$ ($2.2 \times 10^{-3}$) | 0.50 (0.06) | 0.54 (0.04) |
| PrettYharmonize | $2.7 \times 10^{-5}$ ($2.5 \times 10^{-4}$) | $5.4 \times 10^{-23}$ ($2.8 \times 10^{-18}$) | N/A | N/A |
| MIRTH | 0.39 (0.56) | 0.52 (0.58) | 0.47 (0.08) | 0.51 (0.04) |
| Oracle | 0.35 (0.54) | 0.41 (0.55) | 0.47 (0.07) | 0.50 (0.04) |

**Figure 3:**
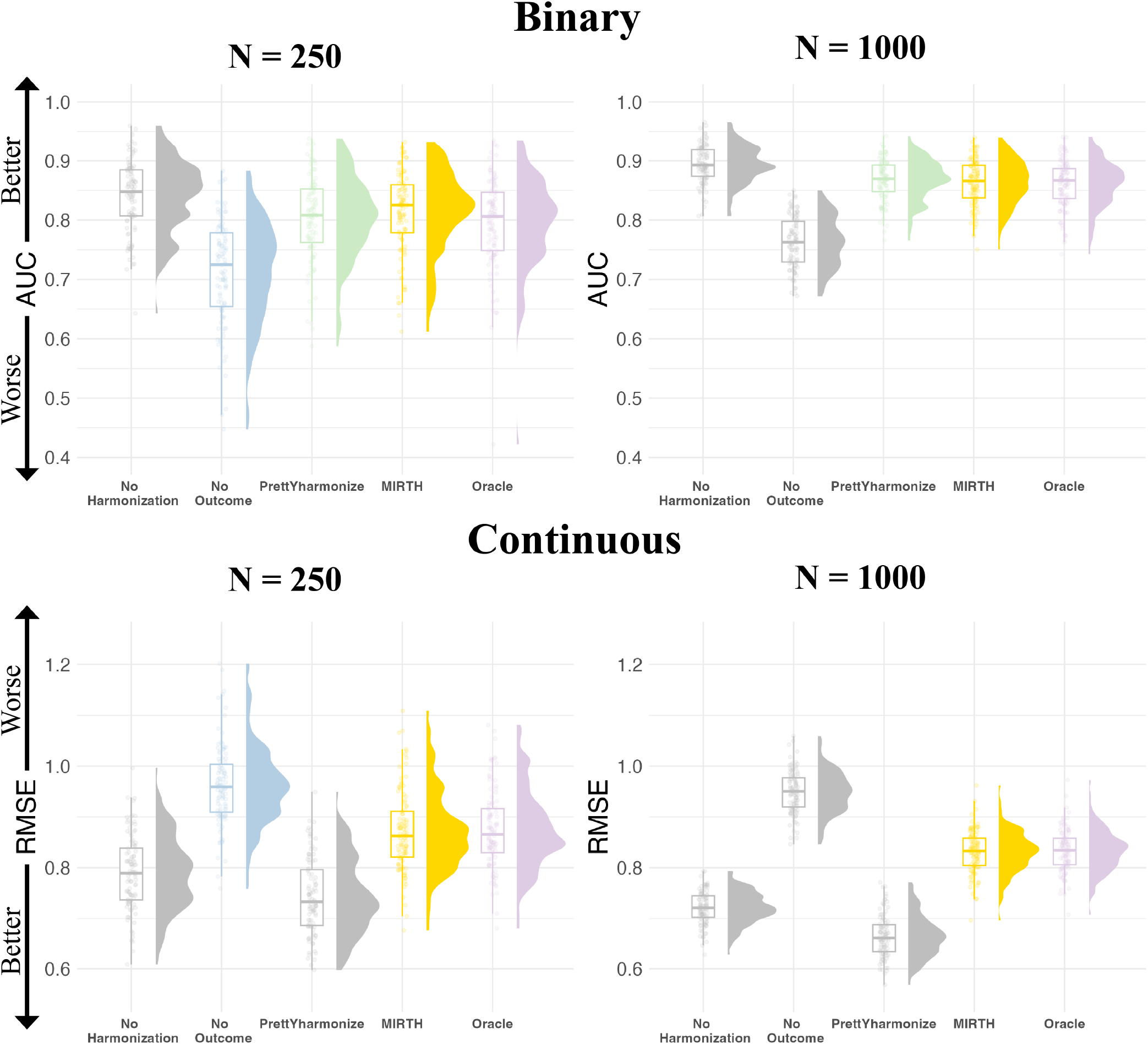
Predictive performance for simulation settings where outcomes are associated with |*V* ^*\**^| = 10 imaging features and *S* = 5. Results depict 100 simulations for each unique combination of sample size and outcome type, with model performance evaluated on a held-out test set (20%). Gray shading indicates that a given method generated predictions that were significantly correlated with site (*p*_median_ *<* 0.05) after controlling for the true outcome, suggesting that site effects may not have been fully removed from the test set (Table 2).

In the last simulation analysis, we tested how MIRTH performed in settings where missingness was present in both the covariate and outcome variable. Table 3 compares results under missingness and no missingness conditions, which exhibited negligible differences in predictive performance across sample size and outcome type (ΔAUC_median_ *<* 0.002; ΔRMSE_median_ *<* 0.003). Thus, MIRTH can readily account for missingness patterns that involve both the outcome and other covariates in the test set.

**Table 3.**
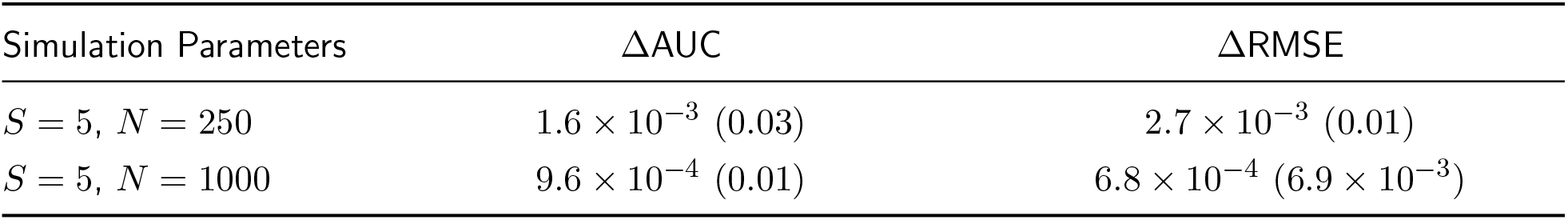
Median (IQR) values across 100 simulations with nonzero imaging-outcome associations, where the difference in predictive performance under covariate missingness is calculated for both binary and continuous simulated outcomes. Section 2.4 contains additional details on the missingness and data generation mechanism.

| Simulation Parameters | $\Delta\text{AUC}$ | $\Delta\text{RMSE}$ |
| --- | --- | --- |
| $S = 5, N = 250$ | $1.6 \times 10^{-3}$ (0.03) | $2.7 \times 10^{-3}$ (0.01) |
| $S = 5, N = 1000$ | $9.6 \times 10^{-4}$ (0.01) | $6.8 \times 10^{-4}$ ($6.9 \times 10^{-3}$) |

### 3.2 Data Analysis

Figure 4 contains results using the ADNI + BLSA volumetric data to predict AD diagnosis. Similar to the binary classification setting in the simulation study, No Harmonization led to the best median AUC across 100 train-test splits (AUC = 0.96), followed closely by PrettYharmonize, MIRTH, and Oracle (AUC = 0.95–0.96), with No Outcome generating the lowest predictive performance (AUC = 0.91). However, when using the harmonized test data to predict site and not AD status as detailed in Table 4, No Harmonization retained site effects (AUC_median_ = 0.83) that were markedly diminished when using other harmonization procedures (AUC_median_ = 0.62–0.64). While the relative predictive performance of each method was generally consistent across evaluation metrics, F1 score (harmonic mean of positive predictive value and sensitivity) was a notable exception. For example, when predicting AD diagnosis the median F1 score for PrettYharmonize was 0.82 — at least 0.08 lower than every other method — even though PrettYharmonize exhibited comparable performance across other evaluation criteria.

**Table 4.** Median (IQR) values across 100 random train-test splits (80% training) examining performance of random forest models to predict site after harmonization of the test set. Since PrettYharmonize does not propose a unique harmonized data set that aims to be independent of site, the method is not included in the table.

| Method | Site Predictions (AUC) |  |  |
| --- | --- | --- | --- |
|  | Diagnosis | Age | Sex |
| No Harmonization | 0.83 (0.02) | 0.84 (0.02) | 0.83 (0.02) |
| No Outcome | 0.64 (0.02) | 0.64 (0.03) | 0.64 (0.02) |
| MIRTH | 0.62 (0.03) | 0.62 (0.03) | 0.62 (0.03) |
| Oracle | 0.62 (0.02) | 0.62 (0.03) | 0.62 (0.02) |

**Figure 4:**
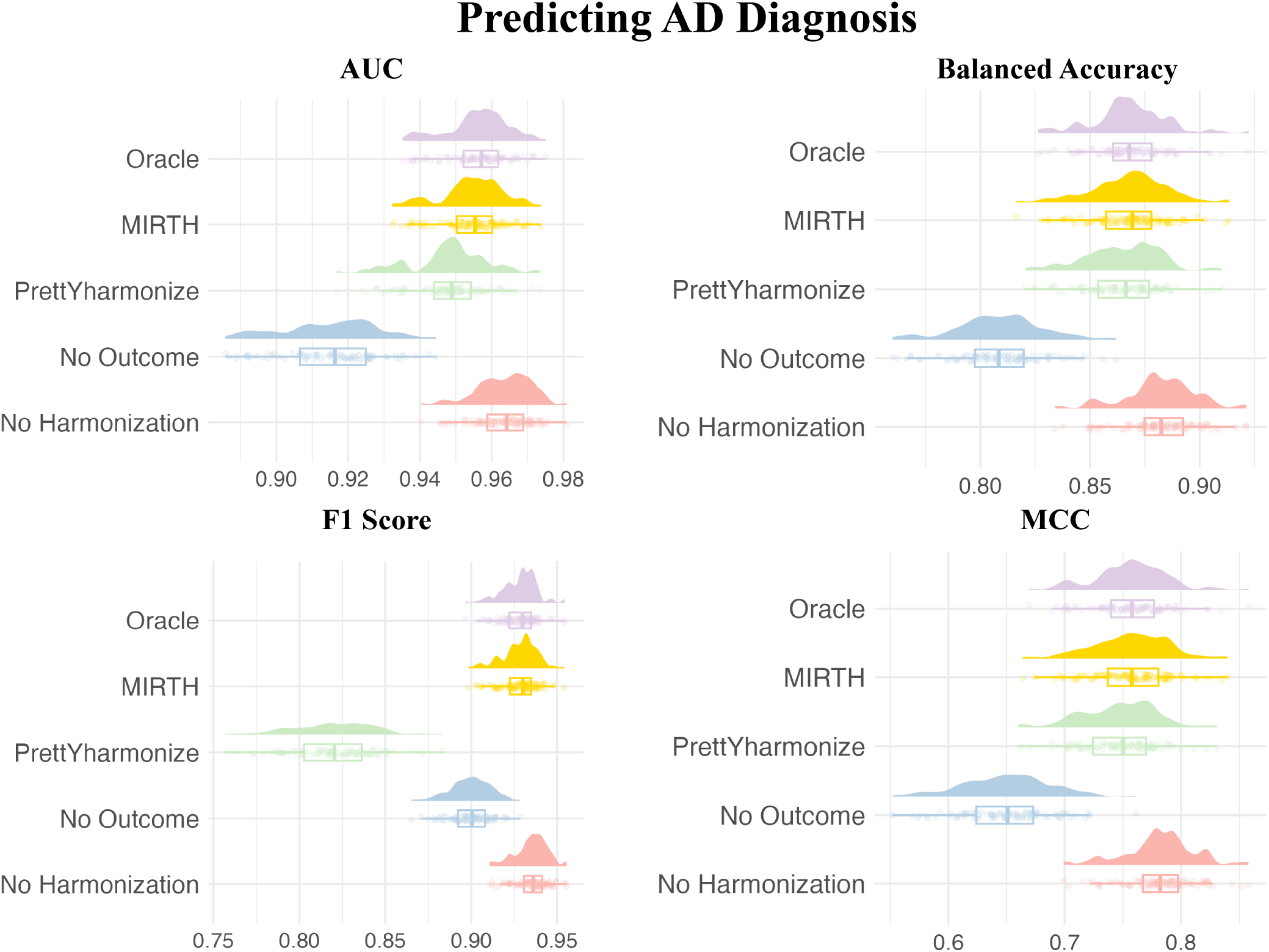
Results depicting predictive performance across four evaluation metrics when predicting AD diagnosis using age, sex, and 145 volumetric features as predictors for a random forest model. Predictive performance is evaluated on the test set (20%) over 100 random train-test splits. Balanced Accuracy 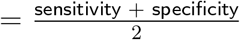; F1 Score =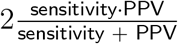 where PPV = positive predictive value, and Matthews Correlation Coefficient (MCC) captures the Pearson correlation between the predicted and actual outcome labels.

To evaluate performance across data types, we also trained random forest models to predict age using the ADNI + BLSA volumetric imaging features. Figure 5 displays predictive performance across 100 random train-test splits for each harmonization procedure and three evaluation metrics. Similar to the binary classification setting, No Harmonization leveraged site effects to inflate predictive performance (RMSE_median_ = 6.88) past the Oracle approach (RMSE_median_ = 7.12). MIRTH (RMSE_median_ = 7.06) exhibited similar performance to Oracle across all evaluation metrics. Consistent with previous results, No Outcome led to poorer predictions due to the attenuation of imaging-age associations during the harmonization process (RMSE_median_ = 7.37).

**Figure 5:**
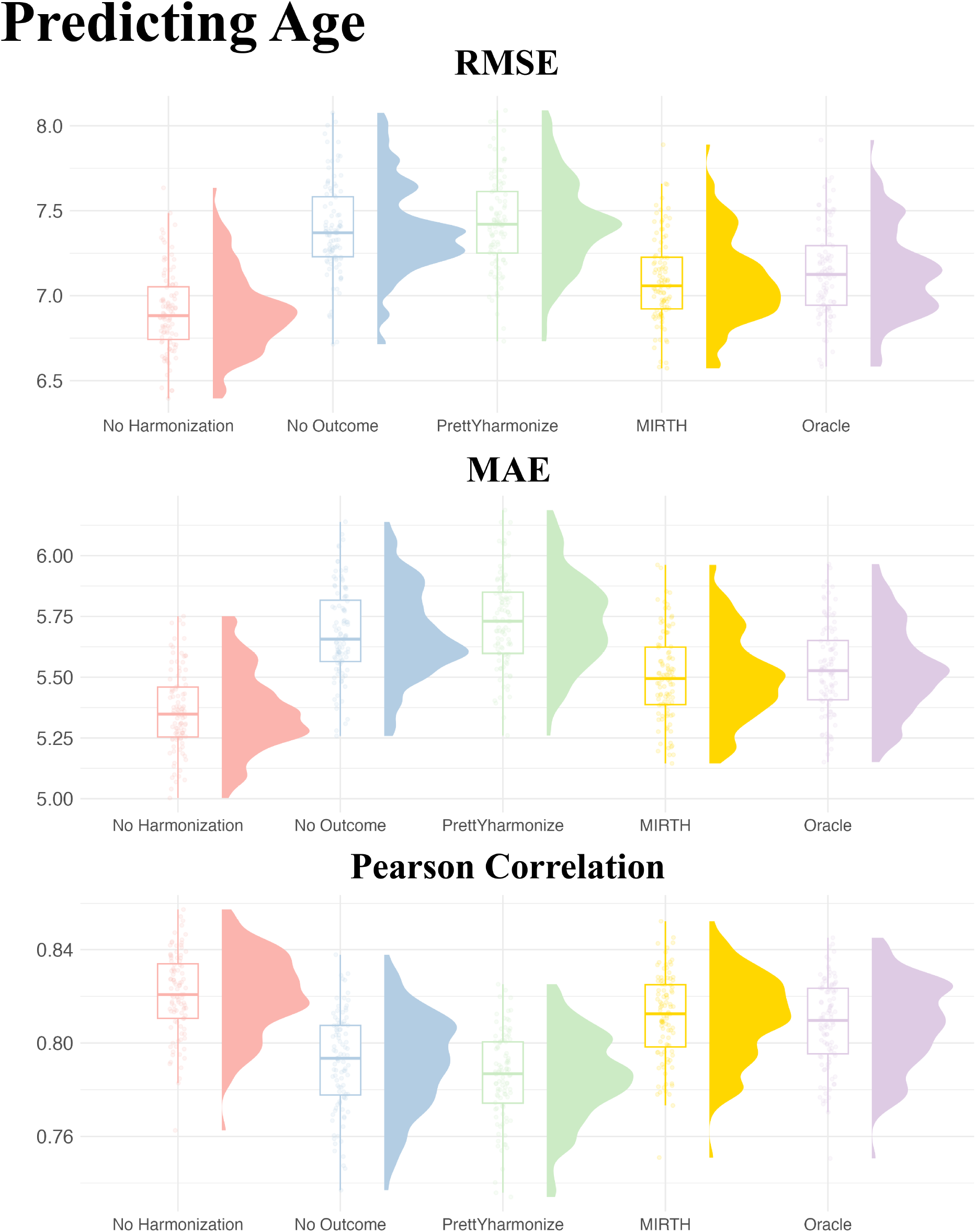
Predictive performance in the test set (20%) across three evaluation metrics and 100 random train-test splits when predicting age using volumetric features, sex, and AD diagnosis. RMSE = root mean squared error and MAE = mean absolute error.

When predicting sex, model performance was consistent across harmonization approaches (AUC_median_ = 0.89). As shown in Figure 6, this result was largely robust to the chosen evaluation metric. However, similar to predicting AD diagnosis, PrettYharmonize exhibited slightly lower F1 scores (F1 score_median_ = 0.80) than other approaches (F1 score_median_ = 0.81–0.82). Consistency in predictive performance across harmonization procedures was likely driven by the outcome being roughly balanced across sites, as shown in Table 1.

**Figure 6:**
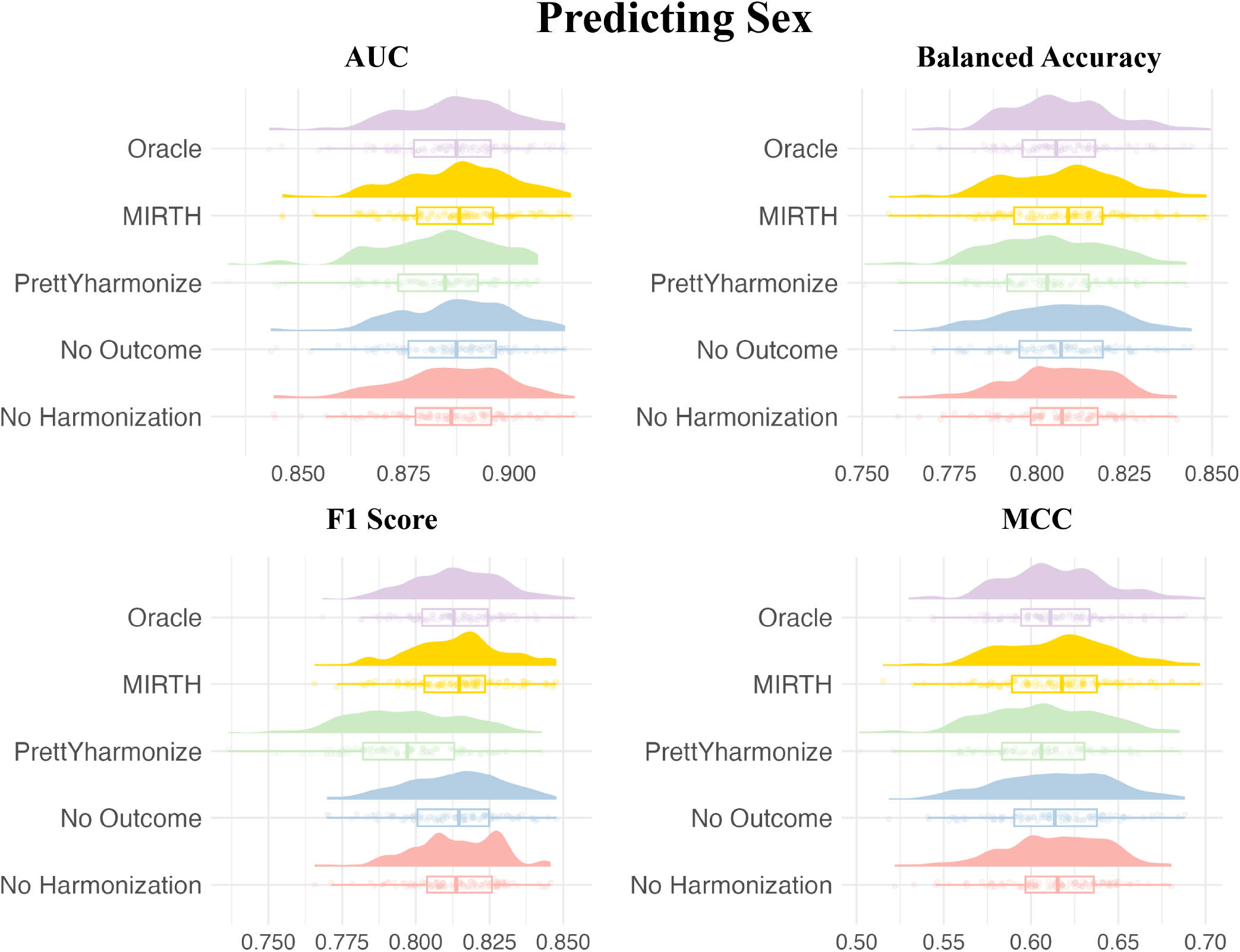
Performance when predicting sex using volumetric features, age, and AD diagnosis as predictors for a random forest model. Model performance is evaluated across four metrics and 100 random train-test splits to ensure robustness.

## 4 Discussion

In this study we proposed MIRTH — a novel procedure for incorporating ComBat into machine-learning pipelines — that generalizes to a broad array of potential missingness patterns in the test data. MIRTH avoids data leakage by fitting the harmonization model on the training data and applying the learned parameters to remove site effects in the test set. To handle missing data, MIRTH utilizes multiple imputation to create several complete test data sets, from which separate predictions can be generated and ultimately pooled. Altogether, MIRTH aims to retain true imaging-outcome associations during the harmonization process via multiple imputation while also avoiding inflated predictive performance due to data leakage or insufficient harmonization of the test data.

In null simulation studies where the outcome was correlated with site but not imaging features, failing to harmonize led to inflated performance, adding to a growing body of evidence that outcome imbalance across sites can markedly impact predictive models if the data are not properly harmonized (Glocker et al., 2019; Nieto et al., 2026; Parker & Leek, 2012; Parker et al., 2014). Moreover, while previous methods may be sensitive to outcome data type, MIRTH’s performance converged towards chance-level as sample size increased for both categorical and continuous outcomes. In simulations where true imaging-outcome associations were present, MIRTH performed comparably to using the true test outcome for harmonization while not producing site-correlated predictions or harmonized test sets that retained technical artifacts sufficient for predicting site.

For simulation settings with missingness in both outcome and covariate variables, MIRTH exhibited comparable performance to when missingness was present in the outcome alone. Thus, current algorithms for multiple imputation such as MICE can be easily adapted to situations where missing data are present across multiple variables, which builds upon previous methods that only consider the missing outcome variable (Nieto et al., 2026). While here we focused on a simple missingness mechanism that multiple imputation can demonstrably address (Murray, 2018; Rubin, 1987), additional work could explore sensitivity analyses for when data are missing not at random.

In a real-world application of predicting AD diagnosis, age, and sex using volumetric features from 145 regions of interest, MIRTH exhibited Oracle-level performance across all outcome variables. When predicting AD/CN diagnosis, predictive accuracy was slightly improved relative to previously reported estimates from the ADNI cohort (Bae et al., 2020; Casanova et al., 2011; Cuingnet et al., 2011; Nanni et al., 2020), likely due to the larger sample size and addition of younger healthy controls from the BLSA study. The No Outcome approach led to noticeably worse performance in this setting, corroborating previous findings that in cases of outcome imbalance across sites, dropping the outcome from the harmonization model can lead to lower classification accuracy (Nieto et al., 2026). For outcomes like biological sex where the prevalence was relatively constant across sites, our results mirrored previous findings (Nieto et al., 2026) that omitting the outcome from the harmonization model has little impact on downstream predictive performance. Thus, whenever the outcome is imbalanced across sites, MIRTH should be strongly considered in order to retain true biological associations during harmonization.

Although MIRTH notably decreased the magnitude of site effects in the test ADNI + BLSA data set, AUC values when predicting site were still above chance (AUC_median_ = 0.62). Visual inspection of variable importance measures revealed that imaging features that predicted site were largely regions where disease effects differed across sites, which violates an assumption of the ComBat model (Jodoin et al., 2025). While MIRTH did not exacerbate this issue, downstream methods that account for disease effect heterogeneity across clinical sites could be incorporated into MIRTH to improve site-effect removal.

This study has some limitations that could motivate future work. First, we focused on incorporating the ComBat model into common machine-learning pipelines. However, recent research has focused on methods that extend ComBat to more complex settings where site effects also influence the covariance between imaging features (Chen et al., 2022; Hu et al., 2024; Ren et al., 2026; Zhang et al., 2023), which may be particularly important in multivariate prediction settings. While there is no obvious barrier to employing MIRTH with other harmonization models, MIRTH’s performance under alternative harmonization specifications should still be explored.

Second, we evaluated MIRTH in settings where the same sites appear in the training and test set. However, in real-world applications, new sites may be present in the test data, such that site-effect parameters need to be estimated in both the training and test set (Xin et al., 2025). When batch effects are large for the unseen site, the initial imputation MIRTH performs may be inadequate, such that iterative procedures that alternate between imputation and harmonization could be further examined.

Third, while we evaluated predictive performance using common evaluation criteria, we did not explore how prediction intervals are impacted by the MIRTH algorithm. One advantage of MIRTH is that the multiple imputation procedure captures uncertainty in the imputation process, such that downstream prediction intervals could be generated that account for the missing data. Future work could explore whether the multiple imputation procedure helps with uncertainty quantification in downstream predictions relative to a single imputation approach.

Altogether, MIRTH can improve multivariate prediction analyses by providing a formal procedure for integrating ComBat into common workflows that retains true imaging-outcome associations, prevents data leakage, and promotes model predictions that are independent of site.

## Acknowledgments

BLSA is a prospective cohort study with continuous enrollment that began in 1958. Data from BLSA are available upon request by a proposal submission through the cohort website (https://www.blsa.nih.gov). The BLSA is supported by the Intramural Research Program of the National Institute on Aging, NIH. This manuscript reflects the views of the authors and may not reflect the opinions or views of the NIH. Data used in preparation of this article were obtained from the Alzheimer’s Disease Neuroimaging Initiative (ADNI) database (https://adni.loni.usc.edu). As such, the investigators within the ADNI contributed to the design and implementation of ADNI and/or provided data but did not participate in analysis or writing of this report. A complete listing of ADNI investigators can be found at https://adni.loni.usc.edu/wp-content/uploads/how_to_apply/ADNIAcknowledgement_List.pdf.

This work is funded by R01MH123550 (RTS), R01MH112847 (RTS, TDS), R01MH113550 (TDS), R37MH125829 (TDS), R01EB022573 (TDS), U01AG068057 (CD), R01CA269948 (CD), and U24NS130411 (CD, HS).

## Code Availability

Code is available at https://github.com/Nhillman19/MIRTH.

